## Supplementary figures and images for "Granick Revisited: Synthesizing Evolutionary and Ecological Evidence for the Late Origin of Bacteriochlorophyll via Ghost Lineages and Horizontal Gene Transfer"

### Supplemental Figure 1

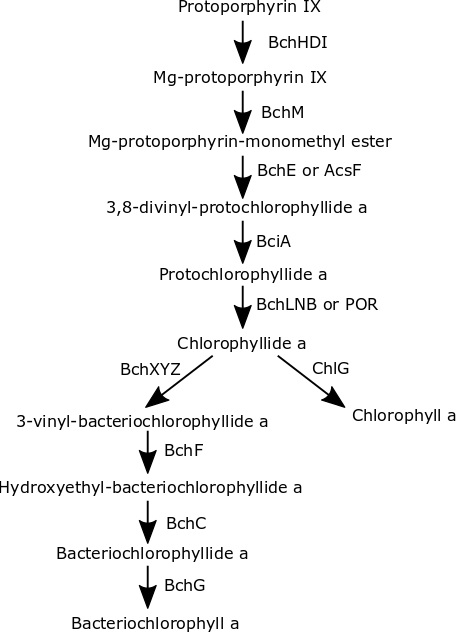

### Supplemental Figure 2a

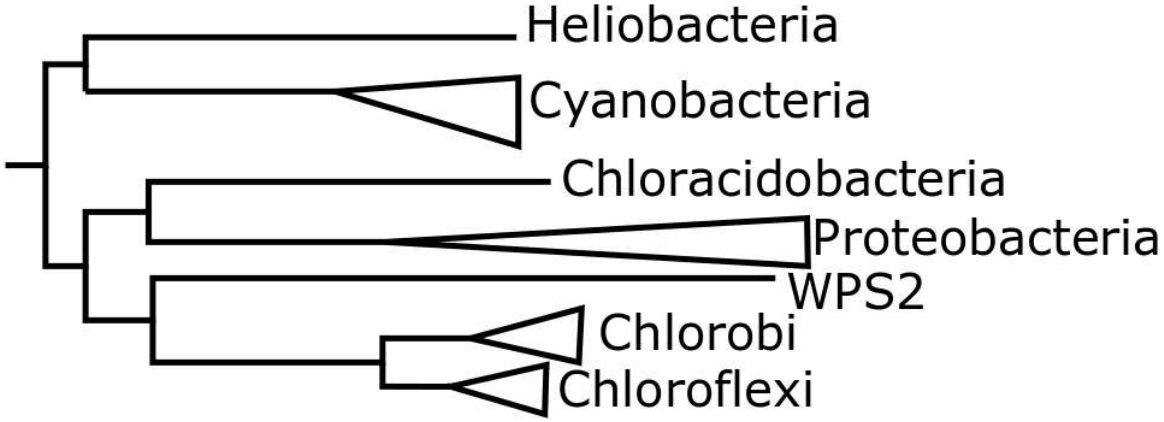

### Supplemental Figure 2b

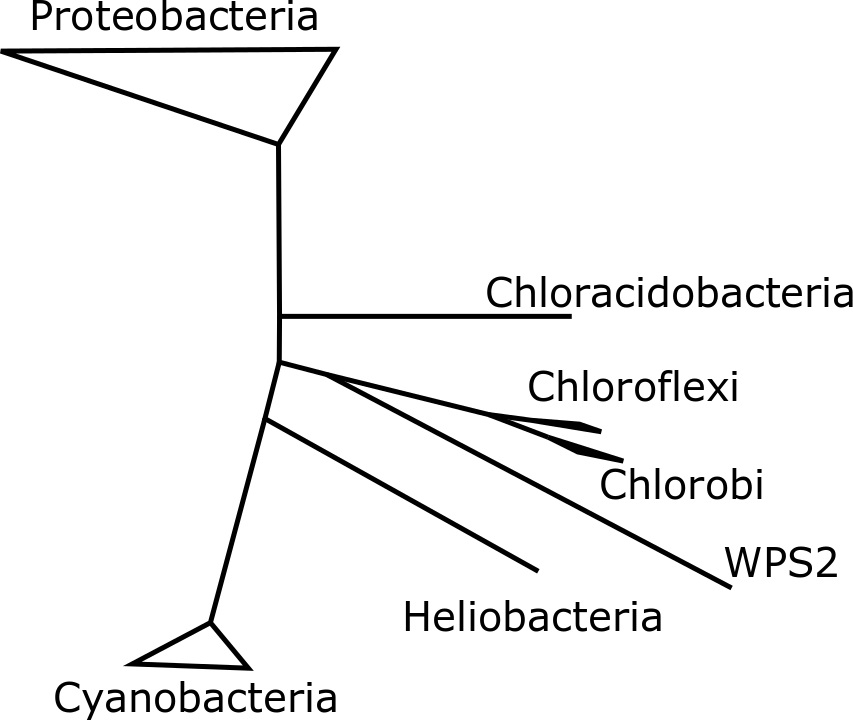

### Supplemental Figure 2c

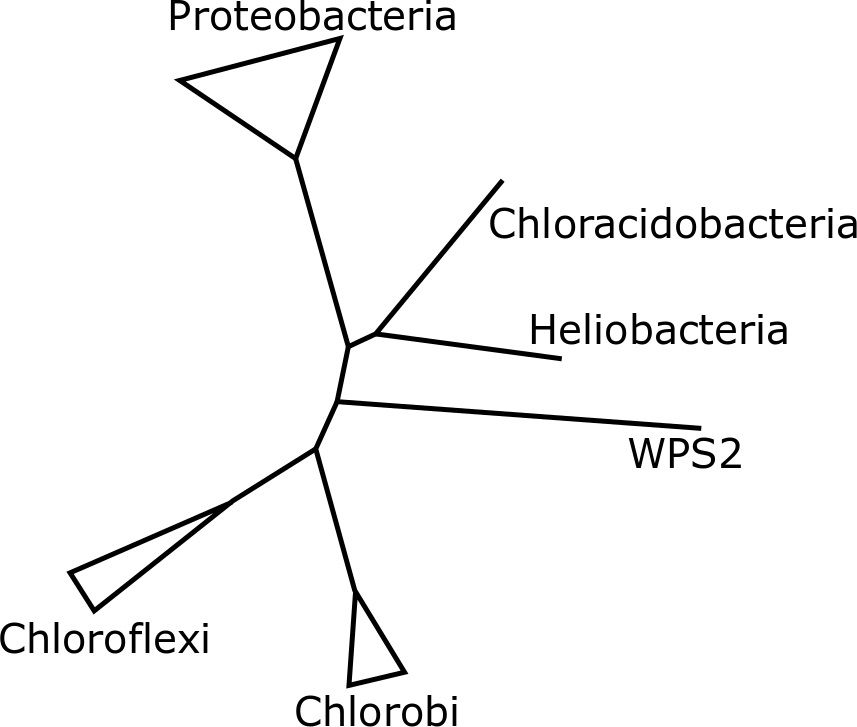

### Supplemental Figure 3

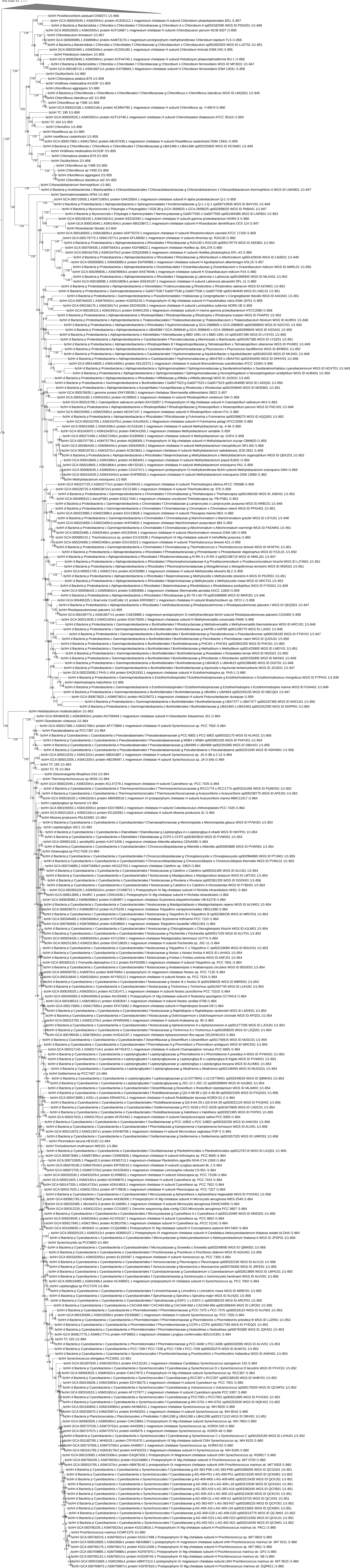

### Supplemental Figure 4

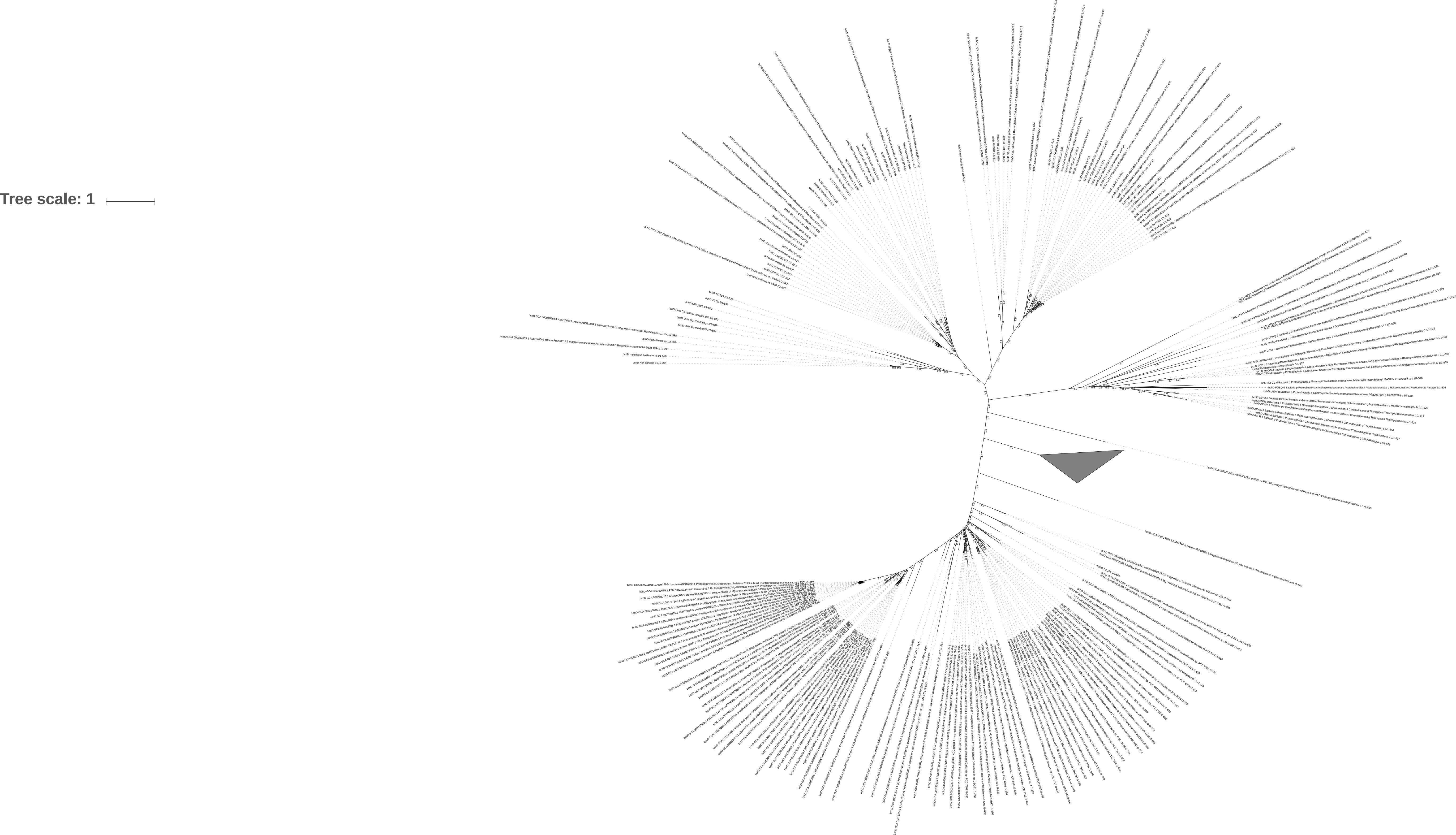

### Supplemental Figure 6

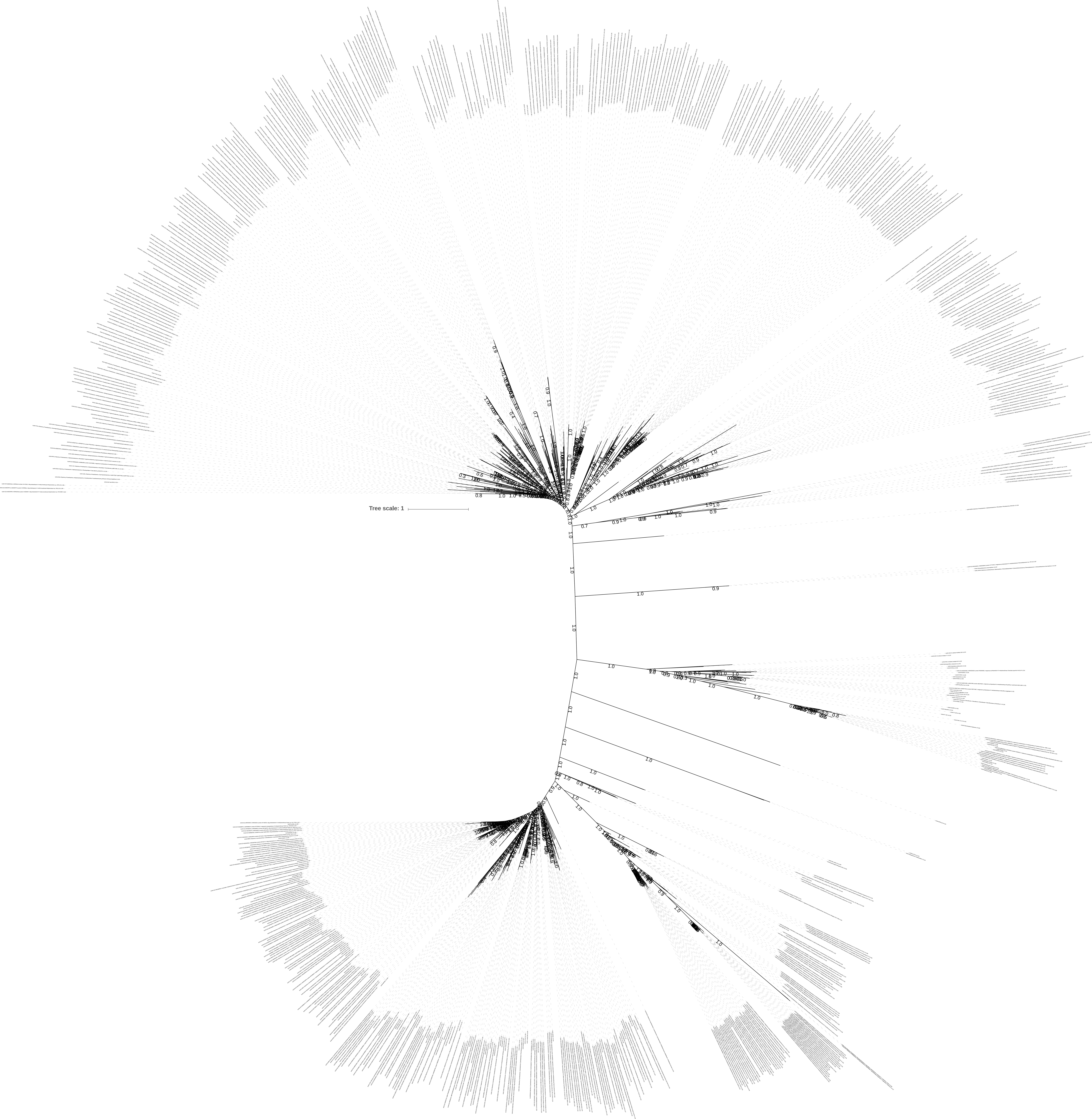

### Supplemental Figure 7

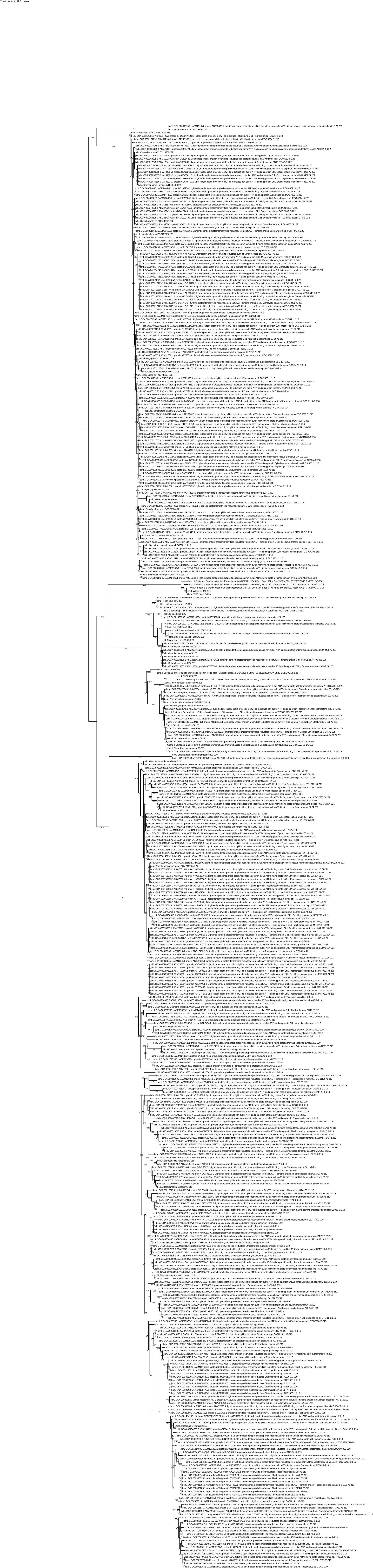

### Supplemental Figure 8

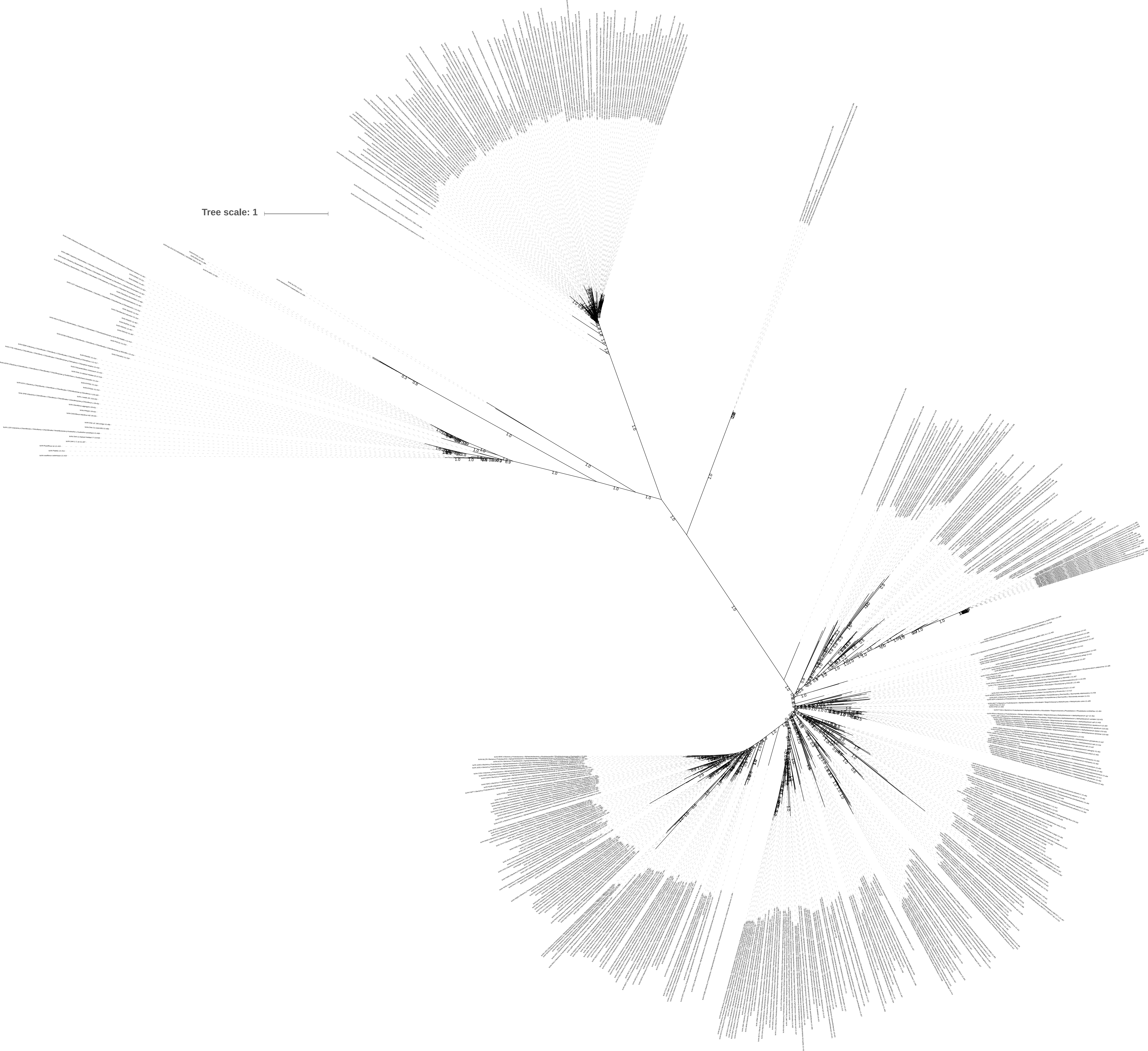

### Supplemental Figure 9

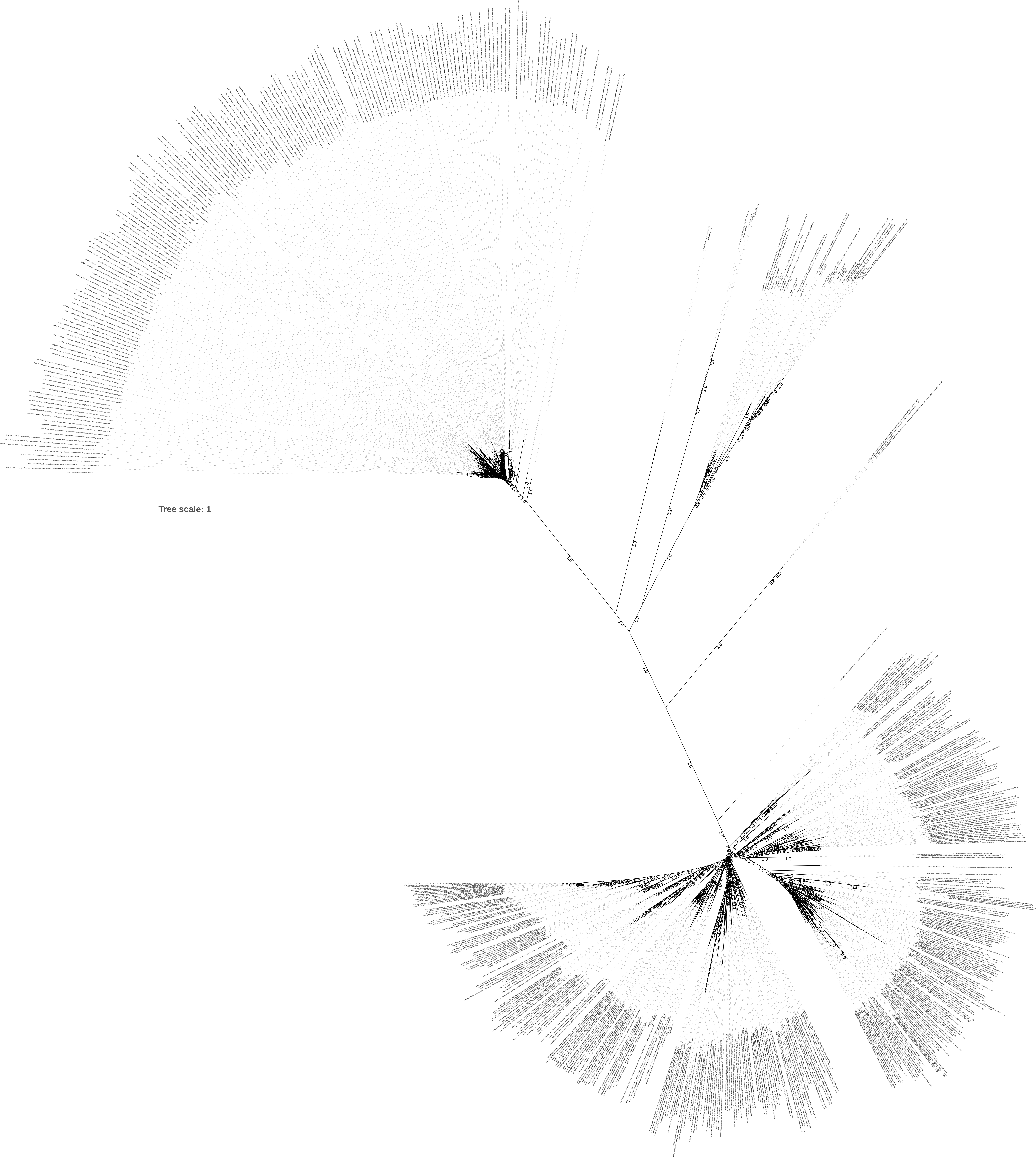

### Supplemental Figure 10

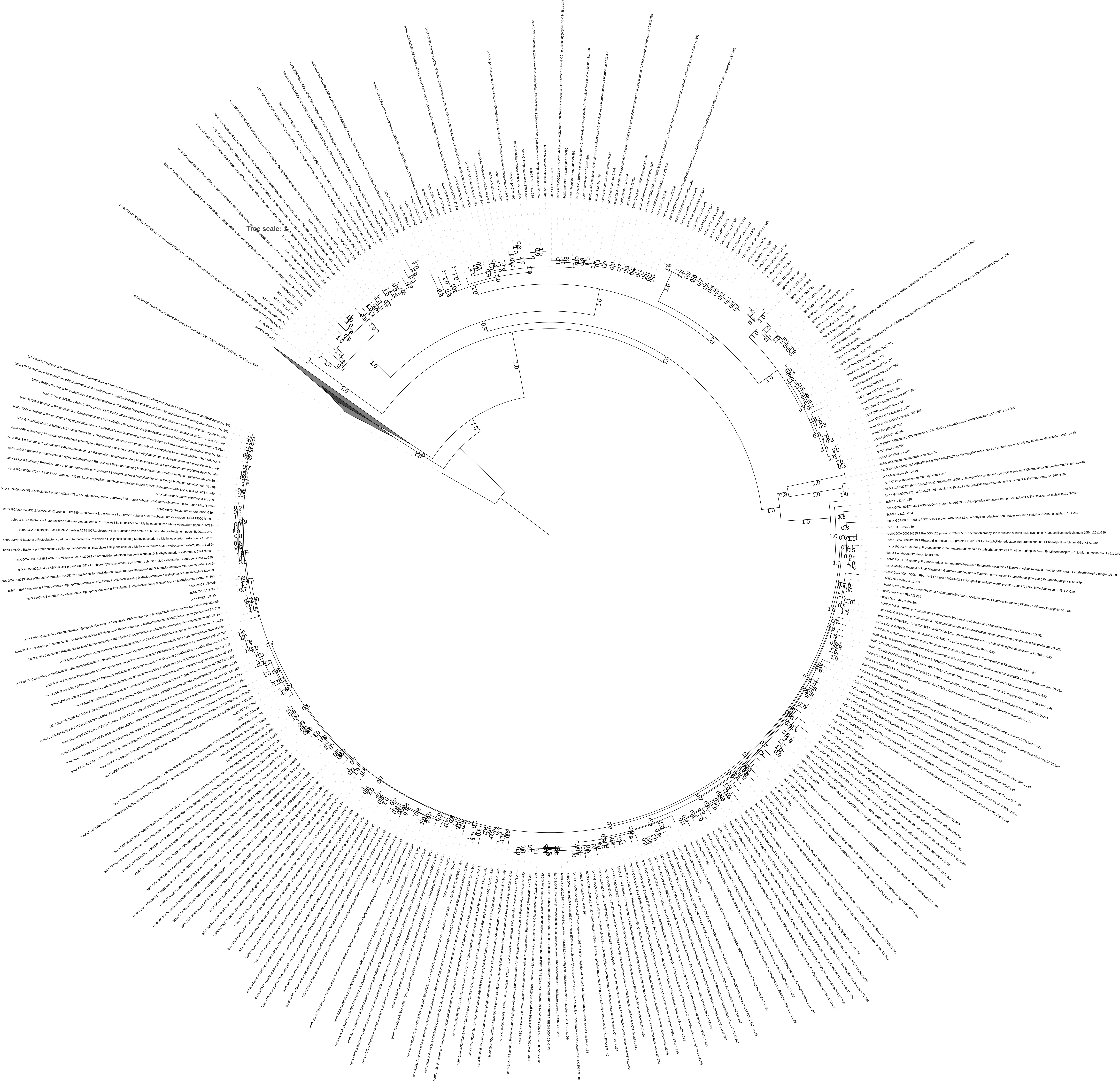

### Supplemental Figure 11

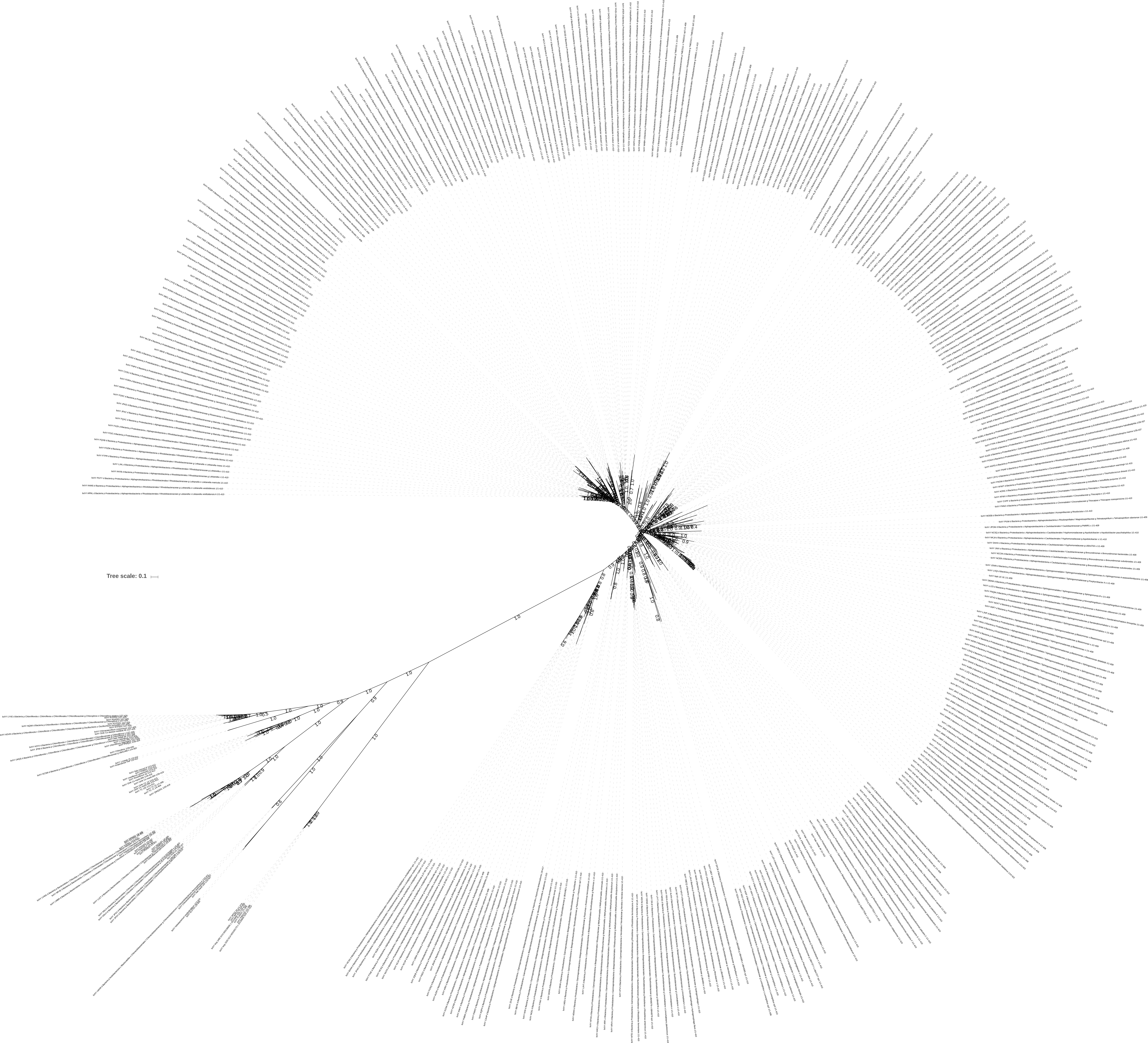

### Supplemental Figure 12

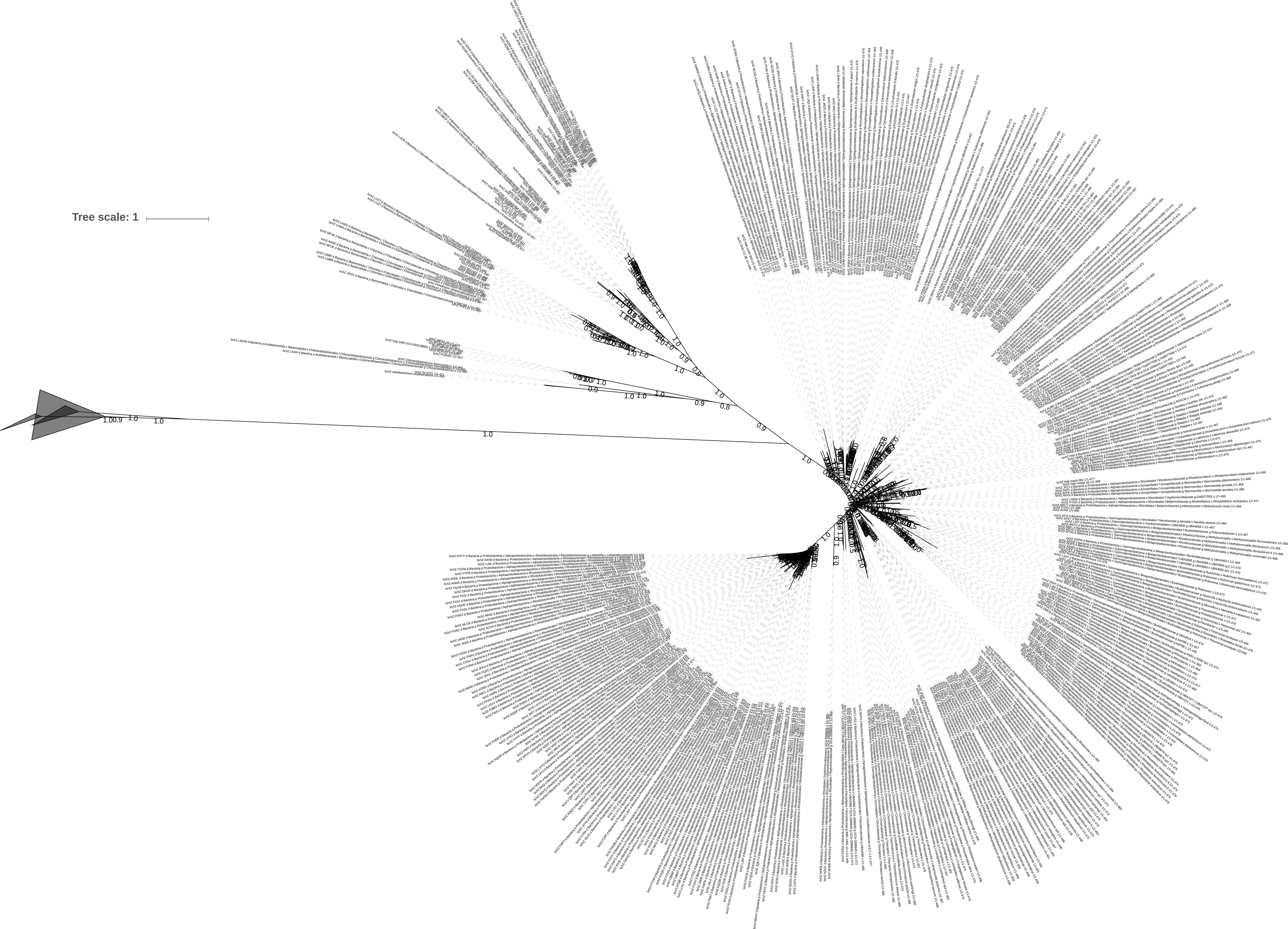

### Supplemental Figure 14

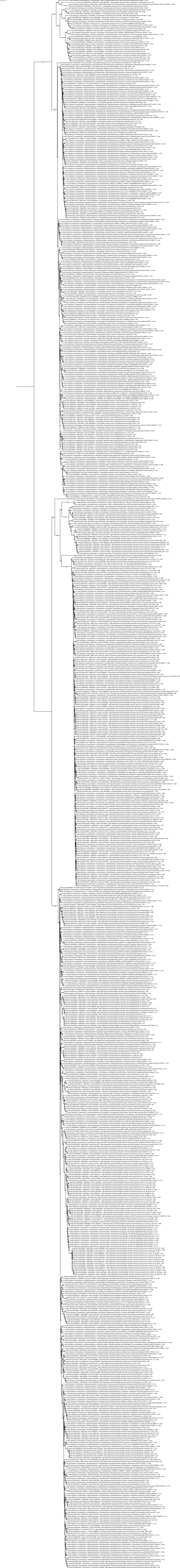

### Supplemental Figure 15

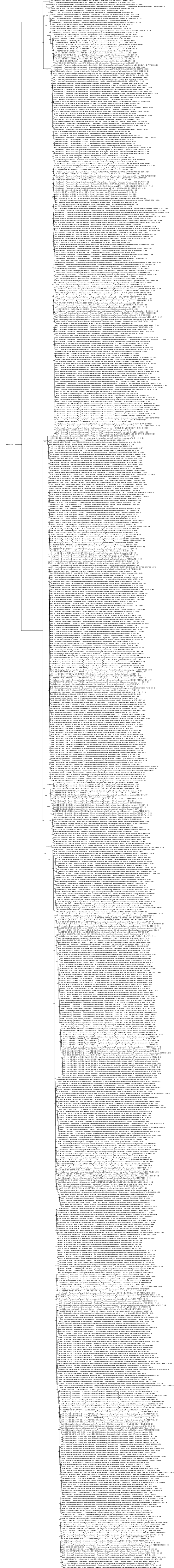

### Supplemental Figure 16

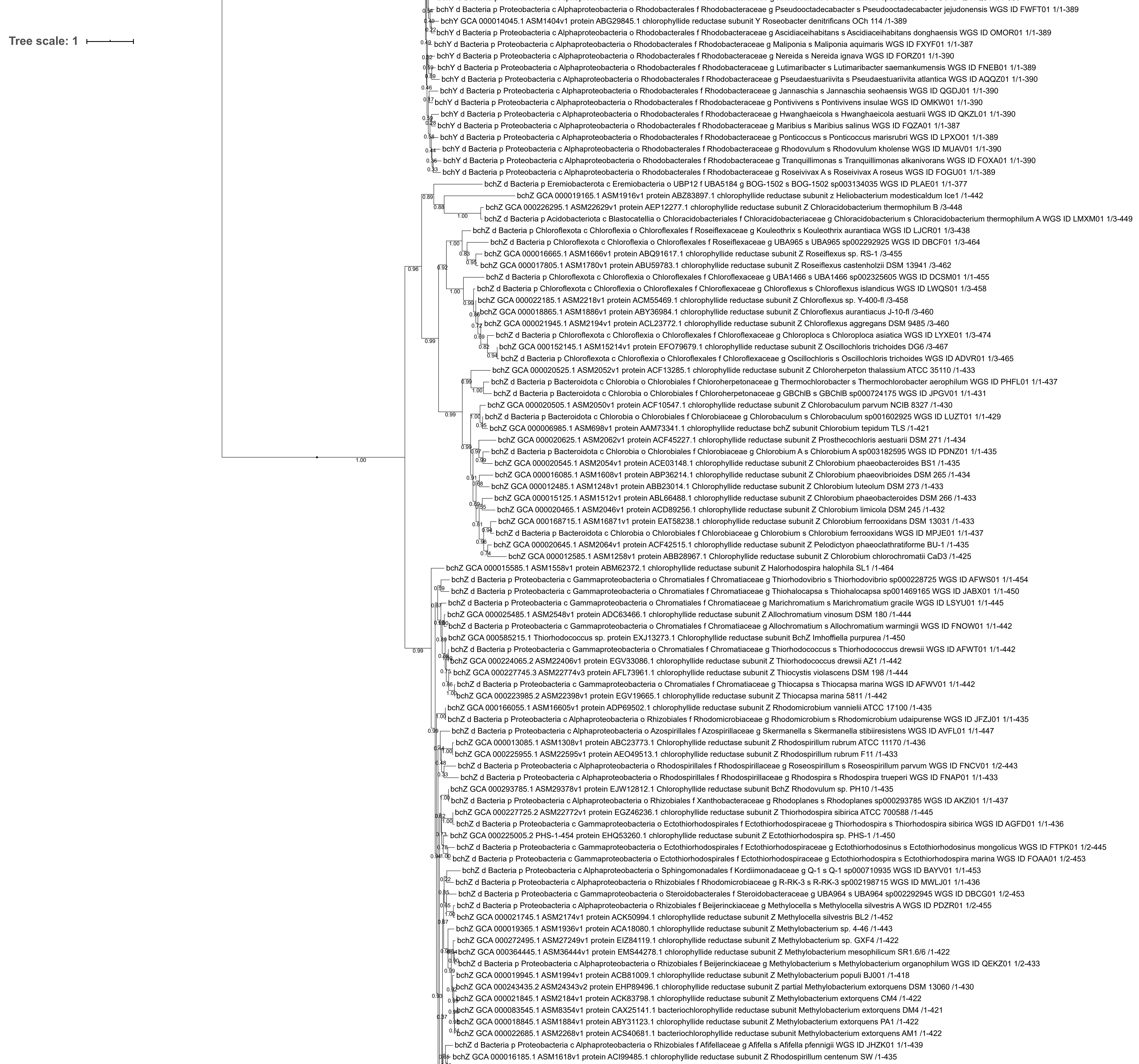
